# Characterizing and comparing phylogenies from their Laplacian spectrum

**DOI:** 10.1101/026476

**Authors:** Eric Lewitus, Hélène Morlon

## Abstract

Phylogenetic trees are central to many areas of biology, ranging from population genetics and epidemiology to microbiology, ecology, and macroevolution. The ability to summarize properties of trees, compare different trees, and identify distinct modes of division within trees is essential to all these research areas. But despite wide-ranging applications, there currently exists no common, comprehensive framework for such analyses. Here we present a graph-theoretical approach that provides such a framework. We show how to construct the spectral density profiles of phylogenetic trees from their Laplacian graphs. Using ultrametric simulated trees as well as non-ultrametric empirical trees, we demonstrate that the spectral density successfully identifies various properties of the trees and clusters them into meaningful groups. Finally, we illustrate how the eigengap can identify modes of division within a given tree. As phylogenetic data continue to accumulate and to be integrated into various areas of the life sciences, we expect that this spectral graph-theoretical framework to phylogenetics will have powerful and long-lasting applications.

---

Phylogenies are essential to many areas of the life sciences. In population genetics and phylogeography, they are used to infer past demography and historical migration events (1). In epidemiology, they are key to understanding how best to control the spread of infectious disease (2). In microbiology, they provide one of the most natural and powerful measures of diversity (3). Phylogenies are also increasingly effective in ecology, where they can inform our understanding of community assembly (4), interspecific interactions (5), and species responses to environmental change (6), as well as guide conservation efforts (7; 8). Finally, phylogenies are essential to comparative phylogenetics (9) and comparative genomics (10) and therefore to our understanding of diversification (11), trait evolution (12), and the genetic underpinnings of both (e.g., (13; 14)).

Despite the importance of phylogenetics in the life sciences, the current techniques aimed at extracting information from phylogenies are limited. One of these techniques is built on summary statistics. In microbiology, ecology, and conservation biology, summary statistics based on measures of phylogenetic diversity, such as total phylogenetic branch length, (7; 15) are often used. In diversification analyses, traditional summary statistics quantify either the stem-to-tip (e.g. *γ* (16) and Lineage-Through-Time plots (17)) or lineage-to-lineage (e.g. *β* and Colless’ index (18)) distribution of branching events across trees. These summary statistics disregard much of the data – and therefore the biological information – encoded in trees: they are simply too crude to precisely capture the complexity of events recorded in empirical trees. Recent computational and conceptual advances based on maximum-likelihood techniques have been able to take better advantage of the full sweep of information provided by empirical trees. Accordingly, they have become the yardstick for determining how clades and traits behave over evolutionary time (9; 11; 19), the selection pressures acting on different genes (20), and changes in rates of infection as a function of time (21). However, all such model-based approaches rely on the a *priori* formulation of a model, which can be problematic, because we cannot exhaustively model the many dynamics potentially generating all empirical trees. Finally, both the summary-statistics and model-based approaches mentioned above are limited to the analysis of ultrametric trees, therefore limiting their domain of applicability. In this paper, we introduce an approach to phylogenetics that does not require any *a priori* assumption about how the phylogeny behaves and can be applied to ultrametric as well as non-ultrametric trees.

We develop an approach based on graph theory that allows a systematic characterization and comparison of the entirety of information encoded in phylogenetic trees. In various configurations, graph theory has been successful in understanding the organizing principles behind biological phenomena at every scale, including the regulation of gene-expression (22), protein-protein interactions (23), metabolic networks (24) and ecological food webs (25). Graph theory and associated spectral analyses have also been useful in phylogenetics, particularly in developing approaches for tree inference (26) or for comparing the phylogenetic composition of microbial samples (27). Metrics like the Robinson-Foulds distance (28) and nearest neighbour interchange (29), too, for example, are used to compare different trees representing the same set of organisms, by counting the number of steps needed to transform one into the other (or both into a third); while others take a geometric approach to define polytopic contours around a reconstructed tree in order to define ‘confidence regions’ in the tree (30). Typically, such distance metrics have been used to identify outliers among or discordance between gene trees, in order to derive a consensus tree or define the ‘space’ that a set of gene trees occupy (31). They are not, however, built (nor adapted) to function as comparative metrics between species trees representing different sets of organisms. Hence, despite the utility of characterizing and comparing phylogenies sampled from different species trees for understanding general principles in the evolution of biological systems, there exists no graph-theoretical approach designed to do so.

The approach we develop here also provides a way to identify distinct modes of division within a tree, which may, for example, reflect distinct modes and/or rates of diversification. Previous attempts in this direction have focused on identifying shifts in diversification rates under a presumed model of diversification. These can, among other things, examine distributions in species richness across the tips of a tree or use other types of imbalance measures (32; 33). More recently, methods such as MEDUSA (34) and BAMM (35) have been developed to detect the location(s) of rate shifts on phylogenies in a likelihood or Bayesian framework, while non-parametric comparisons, based on non-parametric comparisons of branch-length distributions between subclades, identify shifts in rates as well as modes of diversification (36). This latter approach, however, has been implemented only for pairwise comparisons and is therefore not suited for exploring multiple possible modes of division in trees. Furthermore, all above mentioned approaches are limited to the analysis of ultrametric trees.

In the current work, we describe how to construct the spectral density of phylogenetic trees and demonstrate how to interpret this density in terms of specific properties of the trees. We show how to compute the distance between trees based on their spectral densities and how to identify distinct modes of division within individual trees. We use simulations to demonstrate that spectral densities cluster phylogenetic trees into meaningful classes and can identify meaningful modes of division within trees. We illustrate the unique utility of this approach for testing hypotheses on non-ultrametric trees by analyzing different Influenza strains as well as an archaeal tree. Finally, we discuss potential extensions of the approach with implications for the study of community ecology, macroevolution, microbiology, and epidemiology.

## MATERIALS AND METHODS

### Implementation

Throughout, we use the term phylogenetic tree to mean unranked phylogenetic tree shape. Below, we describe how to construct the spectral density profile of a phylogenetic tree, how to compute the spectral distance between trees, and how to cluster trees based on this distance. We also describe how to identify modalities within a given phylogenetic tree and to compute associated support values. We implemented these functionalities in the R package *RPANDA* freely available on CRAN (37).

### Construction of the Spectral Density

Our goal is to provide a common, comprehensive framework for characterising phylogenetic trees, comparing them, and identifying particular branching patterns within trees. We consider a phylogenetic tree as a particular kind of graph, G = (N,E,w), comprised of nodes (N) representing extant and ancestral species, edges (E) delineating the relationships between nodes, and a weight function (w) defining the phylogenetic distances between nodes. We consider fully resolved (i.e., bifurcating) trees throughout for illustrative purposes, but our framework is equally applicable to unresolved trees (i.e., displaying polytomies). We consider trees with explicit branch lengths, but trees with knowledge on only topology could be analyzed using a weight function of 1 for each edge. The framework is equally applicable to ultrametric and non-ultrametric trees, as illustrated below in our empirical applications.

We begin by constructing the modified graph Laplacian (MGL) of a phylogenetic tree, defined as the difference between its degree matrix (the diagonal matrix where diagonal element *i* is the sum of the branch lengths from node *i* to all the other nodes in the phylogeny) and its distance matrix (where element *i, j*) is the branch length between nodes *i* and *j*) (Fig. 1, Supplemental Fig. 1). Each row and column therefore sums to zero. For a tree with *n* tips, there are *N* = 2*n* − 1 nodes, and the MGL is a N x N matrix. The graph Laplacian is said to be ‘modified’ insofar as it takes a distance matrix, rather than an adjacency matrix, as its subtrahend. We also consider a normalized version of the MGL (nMGL), defined as the MGL divided by its degree matrix. The nMGL emphasizes phylogeny shape at the expense of size, which can be useful for comparing phylogenies on considerably different time-scales.

**Figure 1:**
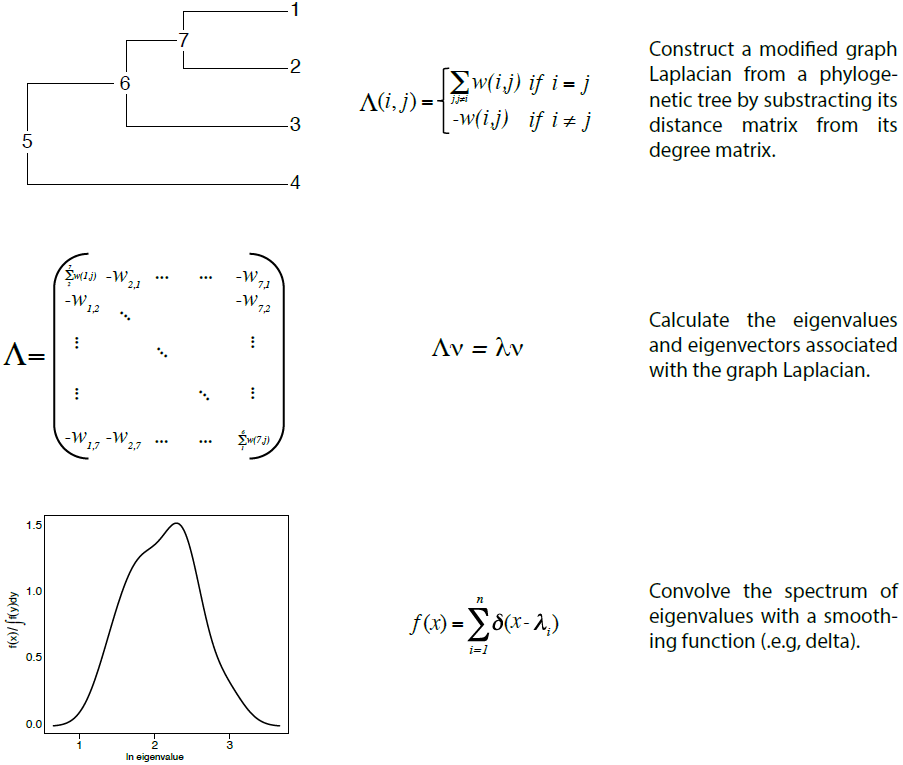
Pipeline for constructing the spectral density of a phylogenetic tree. Graphical depictions (column 1), equations (column 2), and brief descriptions (column 3) for each step in constructing the spectral density are shown. Given a phylogenetic tree with numbered tips and nodes (top left), the Modified Graph Laplacian Λ is computed as the difference between its diagonal degree matrix (where diagonal element *i* is the sum of the branch lengths *w*_*i,j*_ from node *i* to all the other nodes *j* in the phylogeny) and its distance matrix (where element *j*) is defined as the branch length (i.e., the ‘weight’) between nodes *i* and *j*). Next, the eigenvalues *λ* and eigenvectors *v* of Λ are computed (middle row). Finally, the spectral density is obtained by convolving the eigenvalues with a smoothing function (bottom row). See Supplemental Figure 1 for a toy example.

We then construct the spectral density by convolving the spectrum of eigenvalues, *λ*, of the MGL (or nMGL) with a smoothing function. Here, we use a Gaussian kernel,

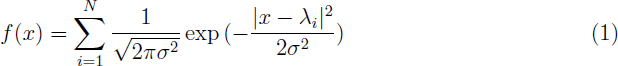

where *N* is the number of *λ* and *σ* = 0.1. The choice of kernel does not considerably change the distribution (38) and the value of *σ* is selected for the degree of desired resolution (i.e., smaller values will highlight finer details at the expense of global ones). The spectral density of a tree is then plotted as a function of *ln*(*λ*) as 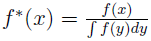. Throughout, spectral densities constructed from the MGL and nMGL are referred to as standard and normalized spectral densities, respectively.

The spectral density can in principle be constructed for trees of any size. However, spectral density profiles of trees with fewer than ~20 tips can be erratic and difficult to compare to larger ones. We therefore discard any trees with fewer than 20 tips.

### Interpreting Spectral Density Profiles

The global distribution of *λ* from a MGL is indicative of the total structure of the tree. Each eigenvector *v*_*i*_ describes a branching event in the tree, and the eigenvalue **λ**_*i*_ associated with *v*_*i*_ describes the inverse diffusion time of the branching event between two nodes (39). Because the diffusion time between two nodes operates principally as a function of the number of branching events between them (40), large *λ* represent short diffusion times characteristic of branching events in speciation-poor regions of the tree (sparse nodes separated by long branches), and small *λ* represent long diffusion times characteristic of speciation-rich regions (dense nodes separated by short branches).

In order to confirm algebraic interpretations of various characteristics of spectral density profiles, we analyzed different spectral properties directly on simulated trees. We simulated birth-death trees with constant speciation (0.2) and extinction (0.05) rates, with 20 time units each, using the R package *TESS* (41). Trees were pruned of extinct lineages and discarded if fewer than 20 lineages survived to the present. A total of 530 trees remained. We constructed the MGL and nMGL and corresponding spectral densities of each tree. The corresponding skewness and kurtosis were computed as 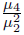 and 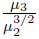, respectively, where *μ*_*i*_ is the ordinary *i*th moment of the distribution. Negative and positive skewness reflect a relative abundance of large and small *λ*, respectively. Lower and higher kurtosis reflect an even and uneven distribution of *λ* values, respectively. We compared the principal *λ*, skewness, and kurtosis of spectral density profiles of each simulated tree to 4 classical measures on these trees: species richness, phylogenetic diversity, the *γ* statistic (16), and Colless’ index (32). Phylogenetic diversity was measured as the sum of phylogenetic branch length (7) using the R package *picante* (42). *γ* is a popular summary statistic reflecting the stem-to-tip structure of a tree: negative *γ* values characterize stemmy trees while positive values characterize tippy trees (16). Colless’ index is a measure of the lineage-to-lineage structure of a tree: smaller Colless’ indices characterize balanced trees, while larger indices characterize imbalanced trees (32). The *γ* statistic and Colless’ index were calculated using the R packages ape and *apTreeshape*, respectively.

### Measuring the Distance Between Spectral Densities

To measure the distance between two phylogenies Λ_1_ and Λ_2_, we begin by computing their spectral densities 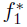 and 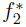, and then use a probability distribution distance metric, the Jensen-Shannon distance, defined as:

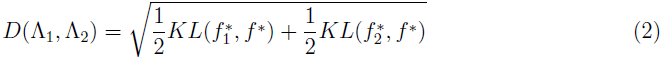

where *D*(Λ_1_, Λ_2_) = *D*(Λ_2_, Λ_1_), *D*(Λ_1_, Λ_2_) = 0 iff Λ_1_ = Λ_1_, 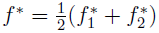, and KL is the Kullback-Leibler divergence measure for the probability distribution, where

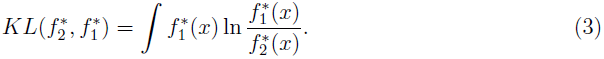

### Clustering Phylogenies from their Spectral Density Profiles

To cluster a given a set of phylogenies, we begin by constructing their respective spectral densities. Next, we compute the Jensen-Shannon distance for each pair. Finally, we cluster the results with energy-based hierarchical and k-medoids clustering, defining the optimal number of clusters by both an expectation-maximization based on the Bayesian Information Criterion *(BIC)* and medoid partitioning. Energy-based hierarchical clustering is a particularly powerful tool for maximizing among-cluster means and minimizing within-cluster means (43) and can show partitioning at different levels of resolution. K-medoids clustering, on the other hand, makes no soft assignments, so each spectral density profile is assigned to a single cluster, and each assignment is given a support estimate based on silhouette width (44).

In order to check the performance of clustering phylogenies based on their spectral density profiles, we implemented the method on a set of trees simulated under different diversification processes using the R package *TESS*. We simulated trees according to six models of diversification, simulating 100 trees under each model, for a total of 600 trees, during 50 time units each. All models had a constant background extinction rate held at *μ* = 0.05. The models had either a (i) constant speciation rate, (ii) decreasing speciation rate, (iii) decreasing speciation rate dipping below *μ*, (iv) increasing speciation rate, or constant speciation rate with an (v) ancient or (vi) recent mass extinction. Speciation rates as a function of time corresponding to each of the four first models and the timing of mass-extinction events corresponding to the last two models are presented in Supplemental Figure 2. Trees were pruned of extinct lineages and any tree with fewer than 20 tips surviving to the present was discarded. We then tested the efficacy of clustering in three ways: using the spectral density profile, using summary statistics of the spectral density profile (principal *λ*, skewness, and kurtosis), and using traditional phylogenetic summary statistics (species richness, phylogenetic diversity, Colless’ index, *γ*, mean branch length, and branch length standard deviation). Spectral density profiles were clustered as described above; summary statistics were normalized and then clustered using hierarchical and k-medoids clustering on principal components.

**Figure 2:**
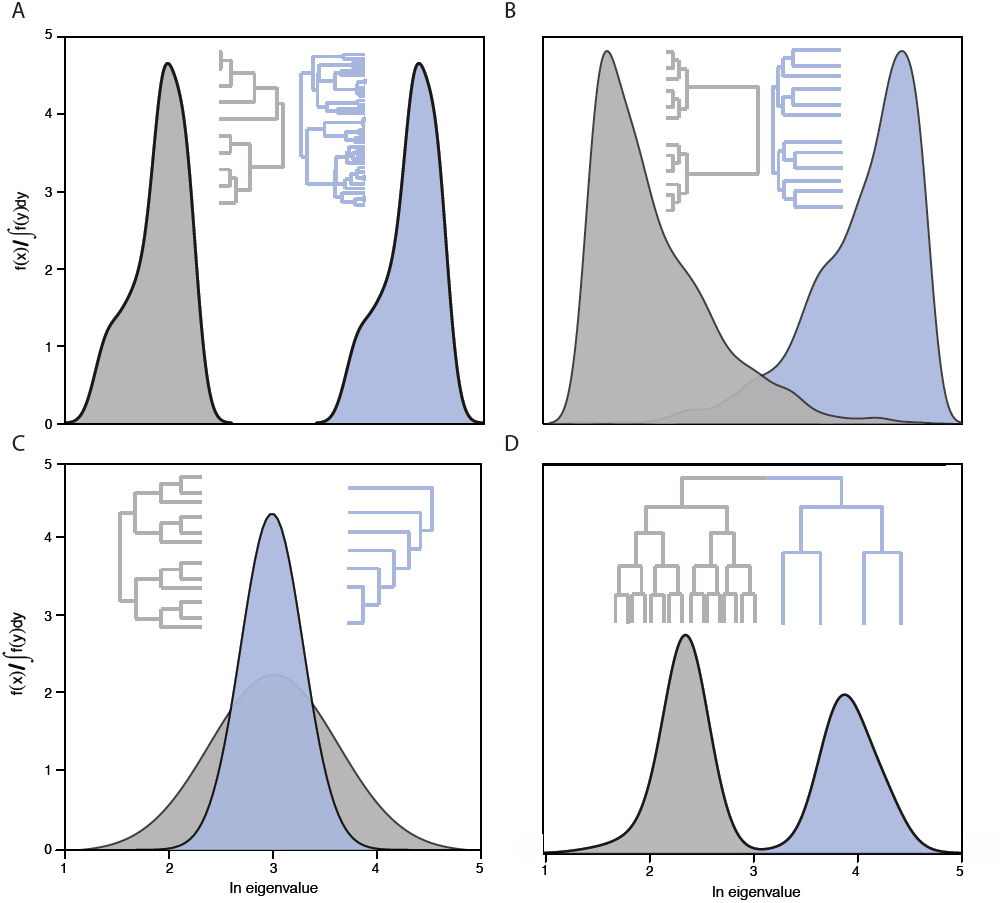
Global properties of the spectral density are indicative of specific patterns in the phylogenetic tree. (A) Trees with high (blue) and low (grey) species richness are characterized by large (blue) and small (grey) principal *λ*, respectively. (B) Stemmy (blue) and tippy (grey) trees are characterized by negative (blue) and positive (grey) skewness. (C) Imbalanced (blue) and balanced (grey) trees are characterized by high (blue) and low (grey) kurtosis. (D) Different modalities within a tree, such as one with stemmy (blue) and one with tippy (grey) branching events, appear as peaks in the spectral density (here at large and small *λ*, respectively).

### Assessing the Sensitivity of Spectral Density Profiles to Undersampling

To assess the effect of undersampling on spectral density profiles, we picked three trees from the simulations detailed above (a constant speciation rate tree, an increasing speciation rate tree, and an ancient mass-extinction tree) and jackknifed (i.e., sampled without replacement) each of them at 90%, 80%, 70%, 60%, 50%, and 40%. One hundred replicate trees were used for each sampling value. We then compared the spectral densities of the complete and undersampled trees, using the Jensen-Shannon distance and spectral properties.

### Identifying Modalities Within a Phylogenetic Tree

To identify modes of division (or modalities) within a phylogenetic tree, we first compute the *λ* from the MGL of the tree and rank them in descending order of magnitude. In graph theory, the ranked *λ* reflect the connectivity of the graph. If there are *i* ideal clusters in the graph (i.e., high among-cluster and low within-cluster variation), then each of the *i* largest *λ* represents a separation between clustered points in the graph. Furthermore, there will be a uniquely large gap between *λ*_*i*_ and *λ*_*i*+1_, where *λ*_>*i*_ << *λ*_≤*i*_ (45). For this reason, the eigengap, identified as the largest difference between two consecutive *λ*, is an indicator of the number of clusters in the graph (40). Transposing this heuristic to a phylogenetic tree, if the eigengap is between *λ*_*i*_ and *λ*_*i*+1_, then there are i clusters, or modes of division, in the tree, and these modalities can be identified using k-medoids clustering on the graph by setting *k* = *i* (46). These clusters need not represent monophyletic regions of the tree, because the *λ*_≤*i*_ describe branching events distributed anywhere in the tree. A cluster could, in principle, be comprised of non-adjacent branches sampled from across the tree.

Once an indication of the number of modalities, *i*, in a tree of interest has been obtained from the eigengap, it is possible to get a confidence measure for *i*. This can be done by comparing BIC values for detecting i modalities in the distance matrix of the tree of interest (*BIC_test_*) and in randomly bifurcating trees parameterized on that tree (*BIC_random_*) (47). Here, *BIC* = *D* + log(*N*) * *m* * *q*, where *D* is the total within-cluster sum of squares based on posterior probability estimates from k-medoids clustering of the nodes of the matrix, *N* is the total number of nodes in the matrix, and *m* and *q* are the dimensions of the clusters of nodes in the x,y-plane, respectively. The random trees can be ultrametric or not. In the former case, tips are randomly coalesced with the same branch length distribution and number of tips as the tree of interest; similarly in the latter case, except the branches are randomly split from the stem. The number of modalities is then considered significant if *BIC*_*random*_/*BIC*_*test*_ ≥ 4 for at least 95% of the random trees. This provides a conservative test of the significance of the modalities. Random trees were constructed using the R package *ape*.

To test this approach, we investigated the ability of the eigengap heuristic to recover simulated shifts in ultrametric trees. We started by simulating a 100-tip pure-birth tree with speciation rate 0.15. A given shift on that tree was simulated by randomly choosing a node located in the middle of the tree (i.e., excluding the first and last quartile of tree length), pruning all branches descending from that node, and grafting onto that node a new tree with proper length simulated under either a pure-birth model with a different speciation rate (ranging from 0.05-0.5) or a different diversification model (randomly chosen among the set of models represented in Supplemental Fig. 2). In a single tree, we simulated up to 10 shifts iteratively, all shifts being comprised of either shifts in speciation rate or diversification pattern (never both). Trees were pruned and grafted using our own code with functions from the R package *phytools* (48). In total, 200 trees with 0-10 shifts in speciation rate and 200 trees with 0-10 shifts in diversification patterns were simulated. The recovery reliability of the eigengap heuristic was compared to MEDUSA (34) and BAMM (35), the most commonly used methods for identifying rate shifts. We ran MEDUSA using the *medusa* function from the R package *geiger* (v2.0.3) with the initial speciation rate set to 0.15 and extinction rate constrained to 0. We ran BAMM after setting priors for each simulated tree with the R package *BAMMtools* (35). We did not compare our results to the non-parametric rate comparison (PRC) of (36), because the approach is implemented only for pairwise comparisons and becomes prohibitively computationally expensive when iterated.

### Empirical applications

To illustrate our approach, we used two empirical datasets: the first one, representing Influenza A strains spanning two animal hosts and 25 countries, was used to illustrate the usefulness of the clustering on the spectral density profile; the second one, composed of 350 16s rRNA archaeal sequences collected from the sediment of Lake Dagow (49), was used to illustrate the identification of modalities within trees. We purposely chose applications on non-ultrametric trees, as analysis of these are not typically available to current techniques.

To put together the viral dataset, we collected trees for a range of Influenza A virus strains from the NCBI Flu Virus Resource (http://www.ncbi.nlm.nih.gov/genomes/FLU/FLU.html). Protein-coding sequences for haemagglutinin (HA), matrix protein 1 (M1), neuraminidase (NA), nucleoprotein (NP), non-structural protein 1 (NS1), polymerase acid protein (PA), and polymerase basic protein 2 (PB2) were obtained for strains from avian and human hosts originating from 25 different countries. Neighbor-joining trees were constructed for each protein separately from multiple sequence alignments in MUSCLE (50). Identical sequences were collapsed. Trees with fewer than 25 or more than 1000 tips were not considered (in a few instances, to meet this criterion, a particular subtype of the strain was selected). In total, 324 trees were constructed. We clustered the standard and normalized spectral density profiles of these trees and compared them with spectral density summary statistics, using peak height as a measure of evenness. (Peak height, defined as the largest y-axis value of the spectral density profile, is a measure of evenness that we found to be better behaved than kurtosis on non-ultrametric trees.) We tested the effect of country of origin on the clustering by randomly assigning strains to clusters and comparing the actual versus randomized distribution of strains for 500 randomizations.

The archaeal phylogenetic tree was simply taken from (49). We removed the four out-groups from the original tree, applied the eigengap heuristic described above to the resulting tree, and characterized each of the identified clusters with their respective spectral density profiles.

## RESULTS

### Constructing the Spectral Density of a Phylogenetic Tree

We construct the MGL of a phylogenetic tree as described above (see *Construction of the Spectral Density*). There is a wealth of knowledge on the graph Laplacian that we may draw from the physical sciences (51). In particular, the graph Laplacian is a positive semidefinite matrix, meaning that it has *N* non-negative eigenvalues, *λ*_1_ ≥ *λ*_2_ ≥ … ≥ *λ*_*N*−1_ ≥ *λ*_*N*_ ≥ 0. Each of these *λ* is associated with a given node and reflects the connectivity of the tree – in terms of both density of nodes and weights – in the vicinity of that node (39). Large *λ* are characteristic of sparse neighborhoods (few nodes) typical of deep branching events, while small *λ* are characteristic of dense neighborhoods (many nodes) typical of shallow branching events (52).

The entire organisation of the tree is best represented as a density profile of the spectrum of *λ* (Fig. 1, Supplemental Figure 1), the so-called spectral density profile (53), obtained by convoluting *λ* with a smoothing function (see *Construction ofthe spectral density*). Importantly, there are heuristic arguments and evidence (although not a formal proof) that it is possible to reconstruct a graph from its spectral density (54), meaning that the spectral density does not lose any of the information contained in the shape of a phylogenetic tree; the only ‘lost’ information is the labeling of the tree. In the physical sciences, spectral density analyses have been successful in differentiating graphs from different domains (55), uncovering network modularity (56), and characterizing synchronization dynamics (57). We therefore hypothesized that the spectral densities of phylogenetic trees would provide powerful tools for characterizing and comparing phylogenies, as well as identifying modules within them.

### Interpreting the Spectral Density of a Phylogenetic Tree

Different aspects of the shape of spectral density profiles may be interpreted in terms of the underlying shape of the phylogeny. In particular, the shift (right bound), asymmetry (skewness), peakedness (kurtosis or peak height), and number of peaks (modalities) of the spectral density profile are illustrative of specific interpretable patterns in the phylogenetic tree. Figure 2 illustrates intuitive interpretations of these four aspects of spectral density profiles. Supplemental Figure 3 demonstrates the validity of these interpretations using trees simulated under a birth-death model with constant speciation and extinction rates (see *Interpreting spectral density profiles*). It also demonstrates that spectral density profile summary statistics and traditional phylogenetic summary statistics are not perfectly correlated, meaning that spectral density profile summary statistics potentially capture different aspects of tree shape.

The *shift* corresponds to the principal (or largest) *λ*, which is related to the largest phylogenetic distance between tip species and may be an indicator of species richness and phylogenetic diversity (Fig. 2A; Supplemental Fig. 3A,B). For the normalized spectral density profile, the principal *λ* is not significantly correlated to species richness and negatively correlated with phylogenetic diversity (Supplemental Fig. 3A,B), demonstrating that the nMGL effectively removes the effect of tree size.

The *asymmetry* of the density profile, which can be quantified by its skewness (a measure based on the 3rd and 2nd moments), is primarily indicative of the stem-to-tip structure of the tree (Fig. 2B; Supplemental Fig. 3C). Intuitively, a positive skewness indicates a relative abundance of small *λ* corresponding to shallow branching events, and therefore characterizes tippy phylogenies, while a negative skewness indicates a relative abundance of large *λ* corresponding to deep branching events, and therefore characterizes stemmy phylogenies.

The *peakedness* of a spectral density profile, which can be quantified by its kurtosis (a measure based on the 4th and 2nd moments), is primarily indicative of the lineage-to-lineage structure of the tree (Fig. 2C; Supplemental Fig. 3D). Intuitively, a flat peak indicates that there is an even distribution of *λ* values, meaning that branch lengths are homogeneously distributed in the tree and so the tree is balanced – while a steep peak, on the other hand, means the tree is imbalanced. Another way to measure this peakedness is by directly measuring peak height. We found that peak height is better behaved than kurtosis on non-ultrametric trees.

The *number of peaks* in the density plot is indicative of the different number of modalities within the tree (Fig. 2D). For example, if clade A is comprised of tippy branching events and its sister clade B is comprised of stemmy branching events, then the spectral density profile of the tree encompassing clades A and B has two peaks, one at small *λ* representing clade A and one at large *λ* representing clade B.

### Comparing and Clustering Phylogenies Using Spectral Density Profiles

Many analyses involving phylogenetics require measuring the distance between phylogenetic trees (an inverse measure of their similarity) and clustering them according to their similarity. Once phylogenies have been transformed into their spectral density profiles, the distance between them can easily be computed using any probability distribution distance metric, and clustered using any traditional clustering algorithm. We used the popular Jensen-Shannon distance metric (58) as our distance metric (see *Measuring the distance between spectral densities*). This metric quantifies the square-root of the total divergence to the average probability distribution; it has the advantage of being symmetric and finite. We used this metric to cluster phylogenies with energy-based divisive clustering, which is a bottom-up hierarchical approach that provides resolution within clusters, and k-medoids clustering, which does not, although it is possible to get support values for cluster assignment using the silhouette width of each data point. Both clustering methods deal well with high-dimensional data (see *Clustering phylogenies from their spectral density profiles*).

In order to investigate the power to identify different types of phylogenetic trees from their spectral density profiles, and to compare this performance to that obtained from traditional summary statistics, we simulated trees under six different models of diversification and checked whether their spectral density profiles clustered into distinct classes of trees. The six models mimicked constant speciation and extinction rates, decreasing or increasing speciation rates (with speciation remaining above or decreasing below extinction rates), and ancient or recent mass-extinction events (Supplemental Fig. 2) (see *Clustering phylogenies from their spectral density profiles*). We found that the spectral density profiles were optimally clustered into six groups with hierarchical clustering (bootstrap probability > 0.95) and k-medoids clustering (P < 0.05), suggesting that spectral density profiles provide an efficient way to distinguish and cluster different types of phylogenies. All models clustered separately with the least and most within-cluster variation in the constant speciation-rate model and the ancient mass-extinction model, respectively (Fig. 3A). A follow-up principal component analysis on summary statistics for the spectral density profile showed comparable influence from principal *λ* (38% of total explained variance), skewness (34%), and kurtosis (28%). Specifically, each acts on a different dimension, with skewness acting orthogonally to principal *λ* (Fig. 3A). Inspection of spectral density profiles representative of each cluster reveals local and global differences between clusters in their distributions of *λ* (Fig. 3B). By comparison, clustering on traditional phylogenetic summary statistics retrieved only three modes of diversification (Supplemental Fig. 4A), which explained 79% of the variance among trees, compared to 93% for spectral density summary statistics. The principal components derived from traditional phylogenetic summary statistics were unable to distinguish between the two decreasing speciation-rate models or between the constant speciation-rate and recent and ancient mass-extinction models (Supplemental Fig. 4B).

**Figure 3:**
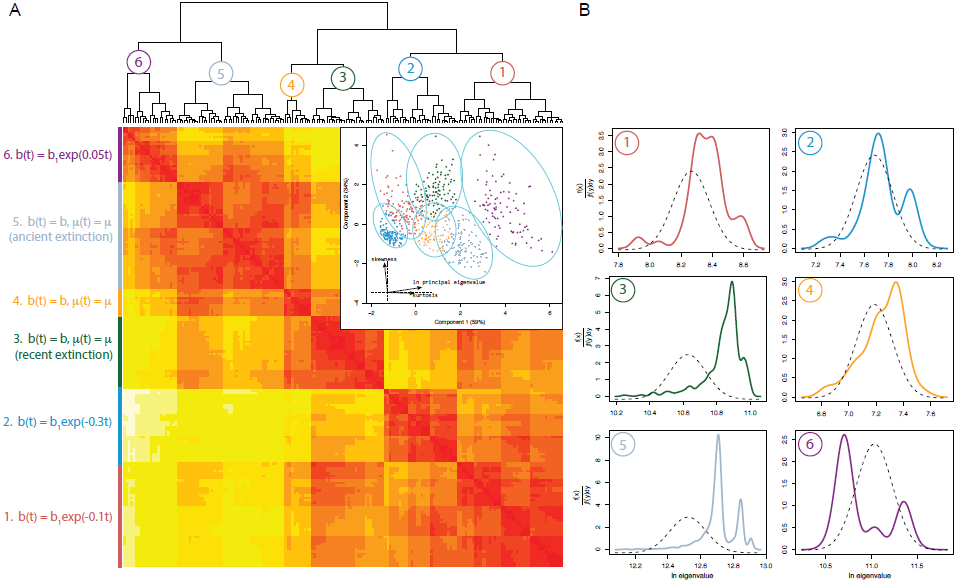
Clustering of spectral density profiles identifies distinct modes of diversification in simulated trees. (A) Hierarchical clustering on the MGL based on the Jensen-Shannon distances between spectral density profiles of trees simulated under different diversification models. Both hierarchical and k-medoids clustering techniques identify six clusters of trees from 600 trees (P < 0.05), each corresponding to a distinct underlying diversification model, whose property in terms of speciation-extinction rate variation is summarized in the left column. Partitions in the hierarchical cluster are collapsed below a threshold height of 2, so that less variation between individual trees is represented as fewer partitions in the hierarchical cluster and fewer cells in the heatmap. (A, inset) K-medoids clustering on principal components derived from spectral density profile summary statistics: ln principal *λ*, skewness, and kurtosis. Shapes correspond to the cluster assignment of trees based on highest silhouette width; tree colors correspond to diversification type. Ellipsoids represent confidence intervals for each cluster, such that each tree could, based on silhouette width support, be assigned to any cluster whose ellipsoid encompasses it. Trees have been assigned to the cluster for which it has the most support. The inset shows the relative contribution of each statistic in the dimensionality of the principal component analysis. (B) A representative spectral density profile for each cluster, defined as the median spectral density profile according to the Jensen-Shannon distance, against a normal distribution (dashed lines) with the same mean and variance, but a fixed height (2.5) to emphasize differences between spectral density profiles, is shown for the six groups. Note the different x- and y-axis ranges.

In order to further test whether phylogeny size was primarily responsible for clustering together the different models, we clustered the same trees using spectral density profiles computed from their nMGLs. We found that these spectral density profiles also clustered by model (Supplemental Fig. 5A), suggesting that trees under a magnitude size-difference are not clustered on size alone. K-medoids clustering on principal components derived from summary statistics calculated on the nMGL, however, retrieved only four clusters, showing an inability to distinguish between the two decreasing speciation-rate models or the constant speciation-rate and recent mass-extinction models (Supplemental Fig. 5B).

### Testing the effect of undersampling on the spectral density profile

Because trees are often incomplete, undersampling is a common issue to consider in phylogenetic analyses. We tested the extent to which (and how) undersampling modifies the shape of spectral density profiles by jackknifing simulated trees (see *Assessing the sensitivity of spectral density profiles to undersampling*). As expected, the spectral density of a tree is sensitive to undersampling and begins to become visually misrepresentative of the complete tree at ~80% complete, although many features of the plot may persist until ~40% (Supplemental Fig. 6A-C). The spectral distance between original and jackknifed trees increased linearly with the level of undersampling. As the trees became less complete, the skewness decreased in constant speciation-rate (1.11 → 0.52±0.10), increasing speciation-rate (0.84 → 0.46±0.09), and recent mass-extinction (1.87 → 1.31±0.14) trees; as did kurtosis, for constant speciation-rate (-0.04→ -1.28±0.12) and recent mass-extinction (2.03 → 0.94±0.14), but not increasing speciation-rate (-0.70 → -0.78±0.21) trees, which showed a sharp increase in kurtosis in some samples at ≤ 50% complete (Supplemental Fig. 8D,E). So, according to their spectral density profiles, undersampled trees are increasingly stemmy, as expected, and, in general, increasingly balanced.

### Identifying Modalities Within Phylogenies

The visual identification of modalities within a phylogeny in the form of peaks in its spectral density profile is only qualitative and not always obvious. A more quantitative approach for identifying such modalities consists in identifying the eigengap, defined as the position of the largest difference between successive *λ* (see *Identifying modalities within a phylogenetic tree*). If the eigengap is between *λ*_*i*_ and *λ*_*i*+1_, then we expect *i* modes of division within the tree. The clusters need not be monophyletic (i.e., there can be clusters within clusters). In some cases, it might be useful to assess the significance of these clusters in comparison to clusters arising by pure stochasticity. This may be done by comparing *BIC* values for finding *i* clusters in the distance matrix of the tree of interest and randomly bifurcating trees (see *Identifying modalities within a phylogenetic tree*).

To assess the ability of the eigengap to recover shifts in diversification, we generated trees with simulated shifts in modes and rates of diversification. We then applied the eigengap heuristic (which includes the post-hoc *BIC* analysis), MEDUSA, and BAMM to those trees and compared the number of recovered *versus* simulated shifts. The eigengap heuristic did not artificially detect shifts when there were none (Supplemental Fig. 7A). The eigengap heuristic and MEDUSA performed comparably well on trees with shifts in only speciation rate (Supplemental Fig. 7A), with both methods routinely underestimating the number of shifts, something previously reported for MEDUSA (35). For trees with shifts in diversification patterns, however, the eigengap heuristic consistently outperformed MEDUSA (Supplemental Fig. 7B). MEDUSA commonly underestimated the number of shifts, while the eigengap heuristic was on average within ±1 the number of shifts. Using the priors estimated from *BAMMtools*, BAMM was not sensitive enough to detect more than three shifts in any tree.

### Empirical Applications

Traditional phylogenetic approaches are typically incapable of dealing with non-ultrametric trees. The ability of our approach to deal with such trees opens up the possibility to analyze the diversification of groups that are rarely studied in macroevolutionary terms. For example, little is known about the diversification patterns of viruses, despite the significance of their evolution for epidemiology (but see (21; 59)). We compared the spectral density profiles of 324 Influenza A trees constructed independently for six proteins, 25 countries, and two animal hosts (see *Empirical Applications*). Results from profiles constructed using the MGL and nMGL showed consistent differences (and similarities) in diversification dynamics across phylogenies derived from different protein segments, originating in different countries, and hosted in different animals. While qualitatively consistent, results from the nMGL were quantitatively more emphatic than those from the MGL, suggesting that phylogenetic shape (not size) was the main effector of differences in diversification. Both showed considerably different profiles for HA and NA compared to the other proteins. Specifically, diversification patterns in HA and NA, in both avian and human hosts, were more expansionary, tippy, and imbalanced than those in the five other protein segments (Supplemental Fig. 8). These differences are demonstrated by a pairwise comparison between HA and PB2 (Fig. 4A-C). We furthermore found, using k-medoids clustering on profiles computed from the MGL and nMGL, six and four (P < 0.05) clusters, respectively, across countries and hosts. Strains from the same country of origin were more likely to fall into the same cluster than expected by chance for avian (D ≥ 0.584, P < 0.001) and human (D ≥ 0.399, P < 0.001) hosts, although this effect decreased when analyzed across hosts (D ≤ 0.185, P ≥ 0.044) (Supplemental Fig. 9). Finally, we found significant differences between hosts for individual strains and across all strains (Fig. 4D-F, Supplemental Fig. 8). We have demonstrated, therefore, how our approach based on the graph Laplacian makes it possible to test macroevolutionary hypotheses on life forms heretofore largely unavailable to diversification analyses; and additionally exemplified how the MGL and nMGL may be used to corroborate different aspects of those tests.

**Figure 4:**
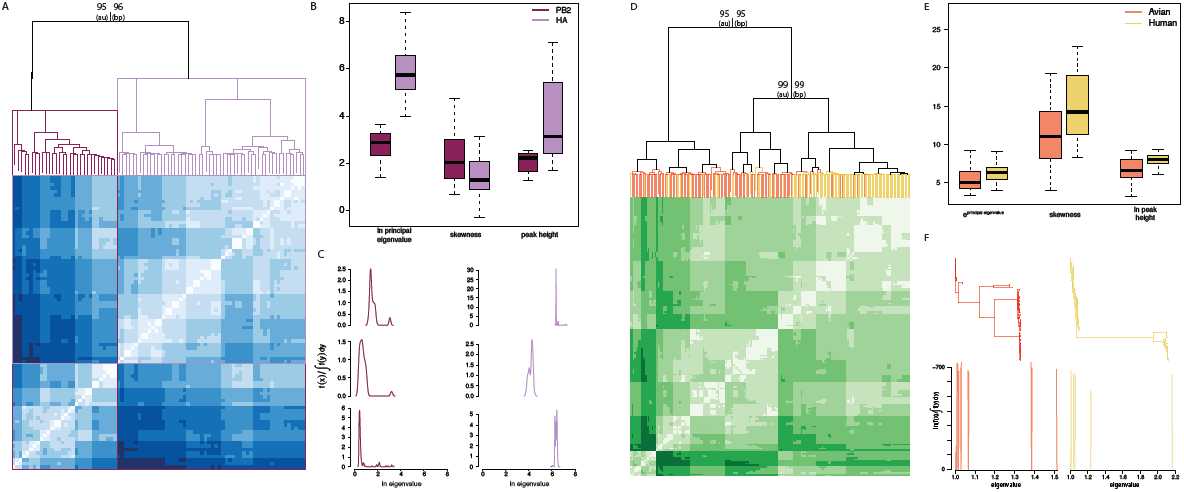
Clustering of Influenza A viral strains from 25 countries and two animal hosts on standard and normalized graph Laplacians. (A) Heatmap and hierarchical cluster of standard spectral density profiles for strains constructed from HA and PB2 sampled across 25 countries in avian and human hosts. Approximately unbiased (au) and bootstrap probability (bp) support are shown at branching events. PB2 phylogenies are overrepresented in cluster 1 (94% of all strains in cluster) and underrepresented in cluster 2 (18%). (B) Boxplot of spectral density profile summary statistics for clusters 1 (light purple) and 2 (dark purple) in (A). All mean differences are significant at P < 0.01. (C) Spectral density profiles from cluster 1 (left) and cluster 2 (right). Note the different y-axis ranges. (D) Heatmap and hierarchical cluster of normalized spectral density profiles for trees constructed from HA, M1, NA, NP, NS2, PA, and PB2 across two hosts and 25 countries. Columns calculated from strains sampled from avian (red) and human (gold) hosts are shown. Phylogenies across all proteins and countries of origin are largely distinguished by host (bootstrap probability > 0.95) (E) Boxplot of mean values for spectral density profile summary statistics. All mean differences are significant at P < 0.05. (F) Trees and spectral density profiles for strains sampled from avian and human hosts.

To further illustrate the empirical applicability of our approach, we examined the spectral density of an archaeal phylogenetic tree of microbial species (see *Empirical Applications*). Using the described framework for finding modes of division within trees, we identified the eigengap to be between *λ*_6_ and *λ*_7_, indicative of six modalities (Fig. 5), which was supported by post-hoc analysis (*BIC_randam_/BIC_archaeal_* ≥ 7.39). Spectral density profiles for the six clusters of nodes (corresponding to the six modalities), as well as where they appear on the archaeal tree, are shown in Figure 5B,C. These results suggest that the archaeal community from Lake Dagow is made of six groups of sequences with distinctive evolutionary histories. The spectral density profile is therefore useful, not only in finding clusters of nodes within trees, but also for assessing what makes these clusters distinct.

**Figure 5:**
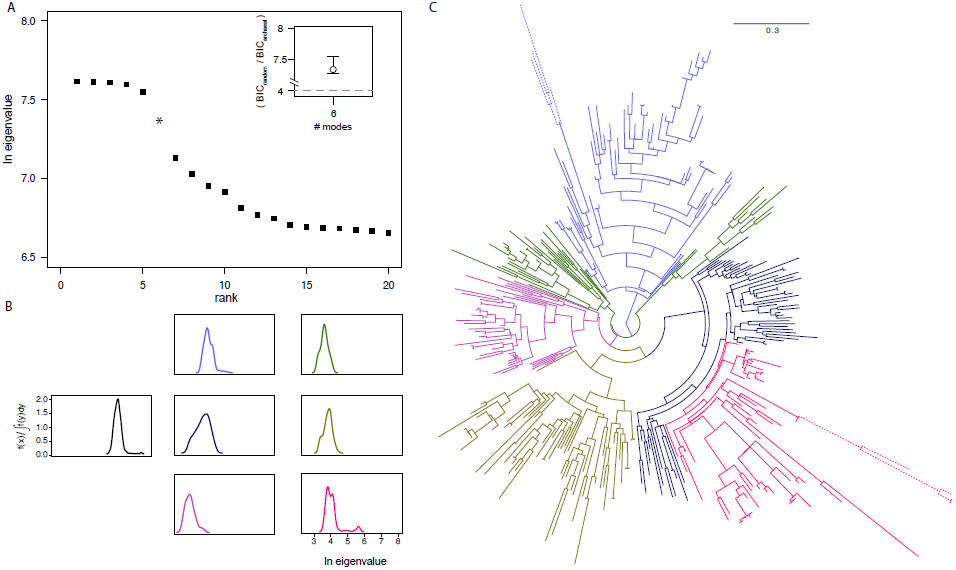
Identifying modes of diversification within a single tree. (A) *λ* calculated from the MGL for the archaeal tree are ranked in descending order and the eigengap is identified between *λ*_6_ and *λ*_7_, suggestive of six modes of division. (A, inset) The ratio of *BIC* values for finding 6 modes in the distance matrices of 100 randomly bifurcating trees and the distance matrix of the archaeal tree. The *BIC* ratio significance cutoff is indicated (grey dashed line). Error bars are drawn from *BIC* ratios against 100 random trees. (B) Spectral density profiles for the original species tree (black) and for each of the modality-trees. (C) Lineages in the archaeal tree showing different modes of division as identified by the eigengap heuristic, with each mode of diversification shown by a different color corresponding to (B). The dashed branches have been shortened for presentation.

## DISCUSSION

We have introduced an approach based on the spectrum of the graph Laplacian for reducing phylogenetic trees to their constituent properties. We have shown how to compute the spectral density profiles of phylogenies, and how to use these profiles to characterize, compare, and cluster trees, as well as to find distinct modes of division within them. This provides a comprehensive framework for 1) summarizing the information contained in phylogenies, 2) identifying similarities and dissimilarities between trees, and 3) picking out distinctive branching patterns within individual trees, without making any *a priori* assumptions about underlying behavior. The ability of this approach to analyze non-ultrametric trees, in particular, fills a largely empty gap in the field of diversification dynamics.

Approaches for summarizing phylogenetic information are required across multiple domains of the life sciences. They are necessary for studying phylogenetic diversity in both the macro- and microbial worlds (3; 7), for measuring how closely related species are within community assemblies (4), for understanding how diversification varies in time and across lineages (11), and for tracking genealogical diversity of infectious diseases through time (21). Such approaches are also particularly useful in modeling approaches, where they allow us to evaluate how closely a specific ecological, epidemiological, or macroevolutionary model reproduces empirical trees. The ability provided by approaches summarizing phylogenetic information to quantify the distance between trees allows us to measure the distance between trees simulated under a specific model and empirical trees, which is crucial to fitting approaches such as Approximate Bayesian Computation (60) or posterior predictive simulations (61).

Given the importance of summarizing the information contained in phylogenetic trees, our study is not the first attempt at doing so. However, our approach is unprecedented insofar as spectral densities account for the entirety of phylogenetic structure, meaning that no shape or size information is lost when expressing a phylogeny as its spectral density. It is therefore superior to previously proposed summary statistics that limit themselves to certain properties of the tree summarized by a single statistic. When reduced to its constituent properties (i.e., principal *λ*, skewness, and kurtosis), the spectral density profile still manages to better identify diversification types among trees than a combination of the most widely used traditional summary statistics. An additional advantage of spectral density profiles compared to many traditional summary statistics is that they can be computed irrespective of whether the tree is dated, ultrametric, or fully resolved.

There are many potential applications of our approach. For example, assuming that co-evolution and co-diversification lead to similarities in branching patterns, clades undergoing co-diversification could be identified based on similarities in their spectral density profiles without any *a priori* information about their interaction. This could be particularly useful in the case of microbes and viruses, for which interactions and co-evolution cannot directly be observed in nature. In viruses, especially, similarities in spectral density profiles can be used to identify convergence across lineages, where diversification may be driven by, for example, an ecological parameter, trait adaptation, or even site-specific substitution. In this respect, our analyses for the various diversification patterns in Influenza A strains – although they are meant here only for illustrative purposes and should be taken with caution – are of some interest.

We find differential effects of protein segment, host, and country of origin on the diversification of Influenza A. For most segments of the virus, diversification patterns are similar, although there are marked differences between both HA and NA and other segments. These two segments show significantly higher mean values for principal *λ* and peak height, indicative of highly expanded, imbalanced diversification, which corroborates previous observations of especially high substitution rates in these proteins (62). Contrary to previous work, however, we do not find similarities in the spectral density profiles of HA and M1, which have been suggested to have comparable phylogenetic histories due to their interaction during viral assembly (63). While these segments may be mechanically interdependent, the considerable variation between their diversification patterns suggests that their strategies of co-evolution, while compatible, are not equivalent. Finally, the exceptional differences between HA and PB2, in particular, with the former exemplifying disproportionately more expansive, imbalanced, and stemmy trees than those constructed with the latter, evinces distinctive evolutionary trajectories for two proteins in a single virus, as well as strong constraints on those trajectories across distant phylogenetic hosts. We furthermore see a significant influence of country of origin on patterns of diversification within each host, where strains from the same country diversify more similarly than expected by chance. However, for both the standard and normalized spectral density profiles, the single strongest impact on the shape of virus diversification is the animal host. These results illustrate the utility of our approach to deal with non-ultrametric trees and to explore the diversification behavior of many organisms previously unavailable to macroevolutionary hypotheses.

Finding shifts in diversification processes is a major interest in macroevolution. Methods for identifying rate shifts in trees (e.g., (34; 35; 36)) have been invaluable in establishing, for example, adaptive radiations in large clades (34; 64). We introduce the eigengap heuristic, an approach for finding different modes of diversification within a single tree. Our approach shows considerable – albeit imperfect – success in recovering rate shifts in simulated trees, comparable (or superior) to the most widely used methods. But it is important to emphasize that the analytic difference in this approach bespeaks a conceptual difference as well: the eigengap heuristic does not strictly identify rate shifts in the tree, but identifies branches of similar diversification processes. So it is not surprising that it underperforms, if only slightly, an existing method in identifying shifts in diversification rate, but outperforms the same method in identifying shifts in diversification pattern. The eigengap heuristic, therefore, distinguishes itself by its power to recognize modes of diversification patterns present in a tree. Our illustration of this approach with an archaeal tree demonstrates how the eigengap heuristic may be used to pinpoint disparately evolving populations of microbial species in a single environment (in this case, Lake Dagow). Specifically, it reveals subtrees with considerably different diversification patterns, which do not vary by phylogenetic relatedness.

Most previous graph-theoretical work in phylogenetics has focused on developing methods to estimate the ‘tree space’ that different hypotheses for the same phylogenetic tree occupy (31; 65; 66). These methods have been very successful and we think that, by assessing the congruence of spectral densities for different gene-based trees for the same species tree, our approach may also be useful for estimating confidence intervals for trees. Similarly, it may be possible to investigate the co-evolution of traits (and genes) based on the (dis)similarities between the spectral density profiles of trait-trees (and gene-trees) sampled from the same species. Generally, comparing spectral density profiles for many phylogenies, whether or not they are sampled from the same species tree, is useful for identifying characteristic patterns of diversification as well as natural limits to those patterns.

There are also many potential variations on our approach. We illustrated the approach on bifurcating trees, yet the degree matrix can take any form, such that reticulated trees (i.e., phylogenetic networks) can also be analysed. Reticulated trees have so far been largely unnavigable by conventional phylogenetic techniques and, as a result, studies of microbial phylogenies have typically assumed the trees to be bifurcating (67), which is often not accurate given the level of lateral gene transfer in the microbial world. Given that microbes constitute the majority of biodiversity on the planet, it is crucial to develop such approaches.

Finally, there are many potential extensions of our approach. For example, graph Laplacians are used in synchronization dynamics (57) to analyze if and how a given part of a network affects the dynamics of other parts of that network. Applied to phylogenies, this could allow for analysing the interaction effects of some clades on others. There are also techniques from differential geometry, based on the so-called trace formula (68), that could be used to analyze the behavior of suites of spectral densities, such as the spectral densities measured for a tree at different times from its origin. Such analyses could inform us about the evolution of a clade. A third potential extension would be to use signed graphs, where a signed matrix maps data onto the edges of the graph Laplacian (69) to analyze how certain information not encoded in the molecular phylogeny (e.g., geographic or phenotypic distance) affects local structures in the tree.

We have developed an approach, implemented in user-friendly software, which will likely become an essential accessory to existing phylogenetic methods by giving researchers access to questions underserved by current phylogenetic techniques.

## Acknowledgements

We would like to thank Julien Clavel, Jonathan Drury, Nancy Irwin, Marc Manceau, and Olivier Missa for helpful comments on the manuscript. EL would like to thank Evan Charles for helpful discussion. Funding was provided by the CNRS and grants ECOEVOBIO-CHEX2011 from the French National Research Agency (ANR) and PANDA from the European Research Council (ERC) attributed to HM.

---

**Supplemental Figure 1:**
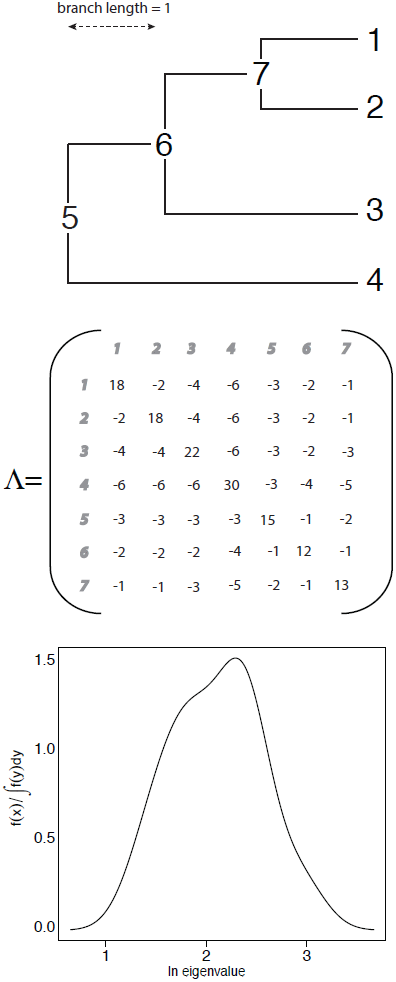
Toy example illustrating the construction of the spectral density for a hypothetical phylogeny. Top: example phylogeny. Middle: computation of the MGL. Each non-diagonal element (*i, j*) in the MGL Λ is equal to the negative of the branch length between nodes *i* and *j*. Each diagonal element *i* is computed as the sum of branch lengths between node *i* and all other nodes *j*. Bottom: the spectral density is obtained by convolving the *λ* calculated from Λ with a smoothing function.

**Supplemental Figure 2:**
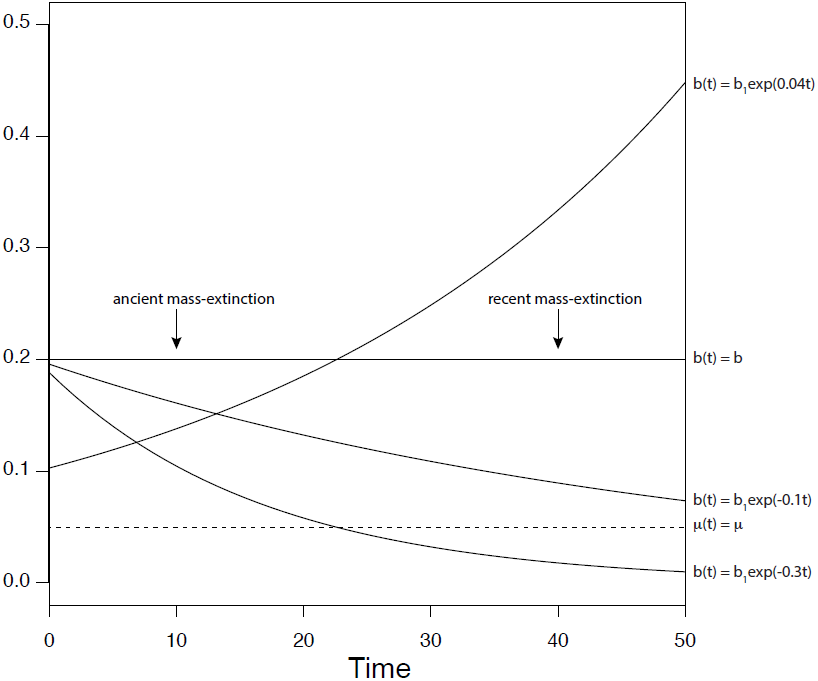
Speciation values for diversification models. Birth-death trees were constructed according to one of six diversification models: increasing speciation, decreasing speciation, decreasing speciation below extinction; and constant speciation-extinction with (i) ancient mass-extinction (0.1 survival probability), (ii) recent mass-extinction (0.1 survival probability), or (iii) no mass-extinction. For all models, *μ* = 0.05.

**Supplemental Figure 3:**
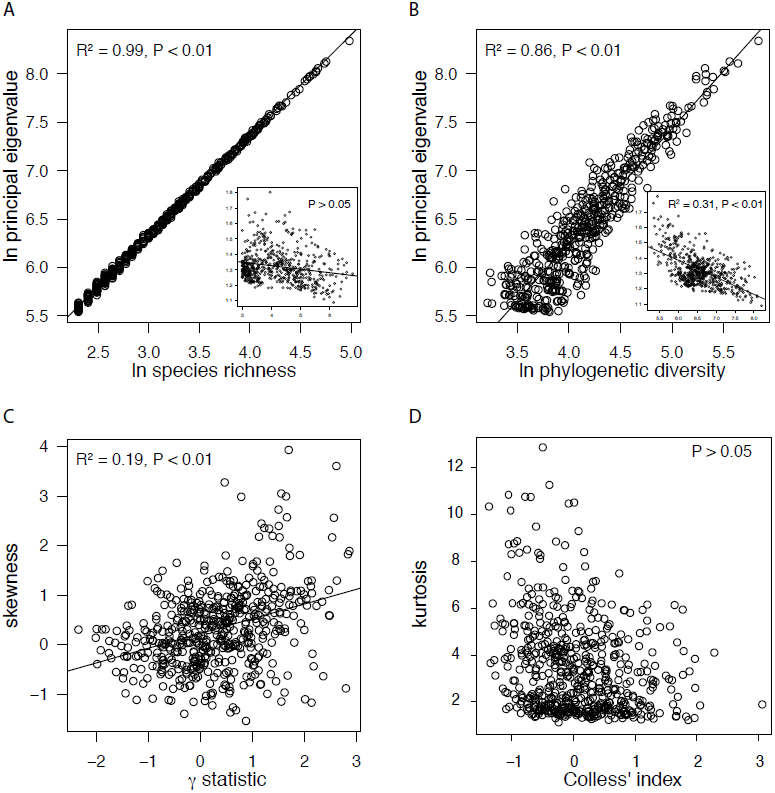
Interpreting spectral density profiles. For trees simulated under a constant birth-death model (open circle), the principal *λ* for the MGL is a good predictor of species richness (A) and phylogenetic diversity (B); there is a significant positive relationship between skewness and the *γ* statistic (C) and no significant relationship between kurtosis and Colless’ index (D). The principal *λ* for the nMGL shows no significant relationship with species richness (A, inset) and a significantly negative relationship with phylogenetic diversity (B, inset). Trees simulated under decreasing (closed circle) and increasing (square) speciation rates, skewness (E,F) and kurtosis (G,H) show different scaling relationships with species richness, phylogenetic diversity, and *γ*. Only significant slopes are shown.

**Supplemental Figure 4:**
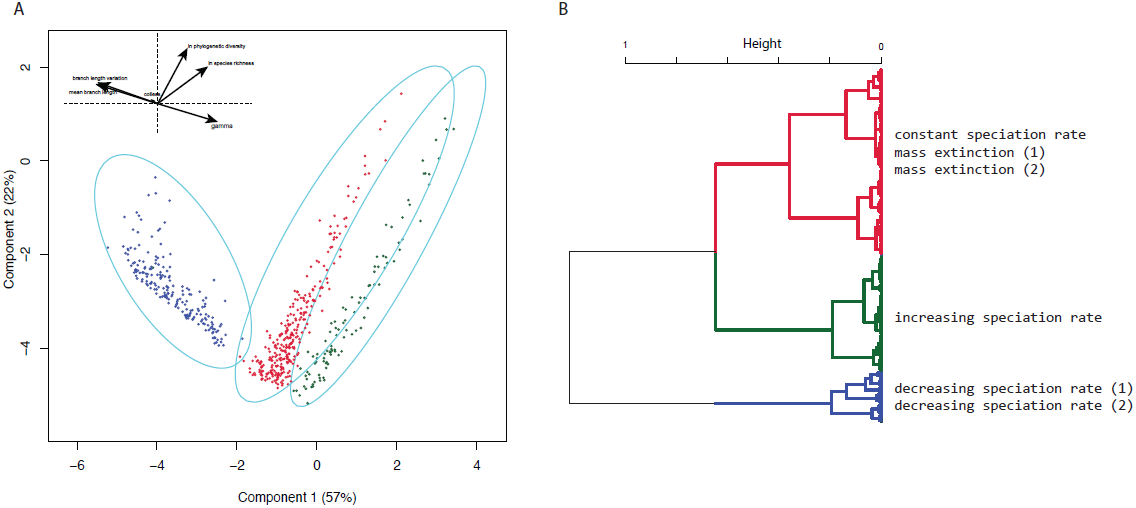
Principal components analysis of simulated trees using traditional summary statistics. (A) K-medoids and (B) hierarchical clustering on principal components derived from mean branch length, branch length standard deviation, ln species richness, ln phylogenetic diversity, Colless’ index, and *γ* calculated for 600 trees simulated under different diversification models. Both hierarchical clustering (bootstrap probability > 0.95) and k-medoids clustering (P < 0.05) extract three clusters of trees. Shape and color correspond to cluster assignment.

**Supplemental Figure 5:**
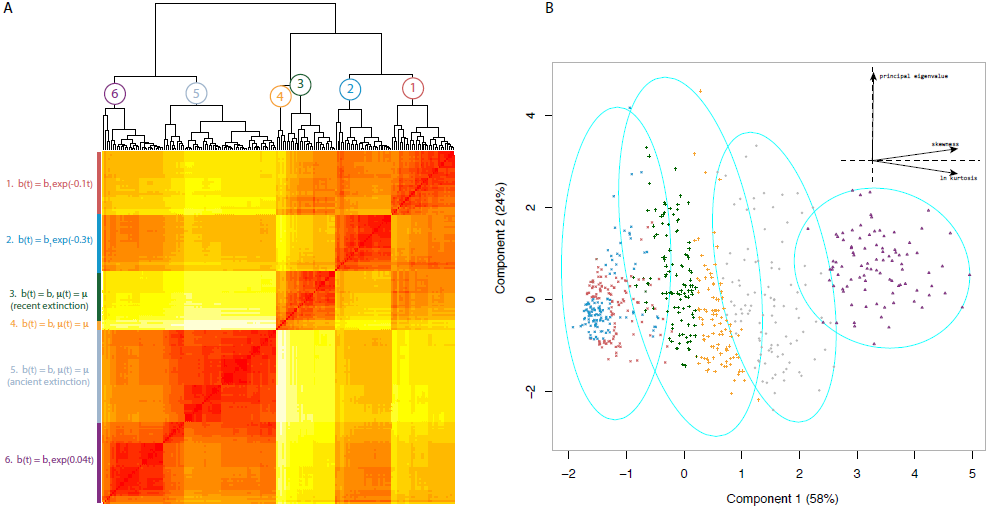
Clustering on principal components and spectral density profiles from the nMGL. (A) Hierarchical clustering on spectral density profiles and (B) k-medoids clustering on principal components are computed as in Figure 3, except based on the nMGL. In (A), hierarchical clustering on the spectral density profile identified six clusters of trees (bootstrap probability > 0.95), each corresponding to a distinct underlying diversification model, whose property in terms of speciation-extinction rate variation is summarized in the left column. In (B), only four significant clusters were identified (P < 0.05).

**Supplemental Figure 6:**
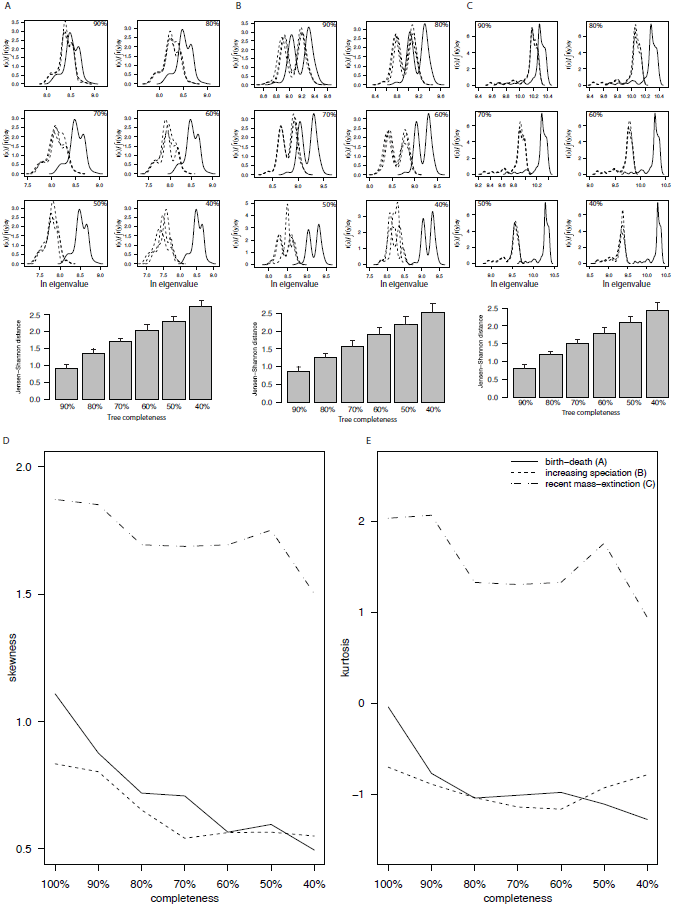
Undersampling affects spectral density profiles. The spectral densities of 3 (out of 100) trees (solid line) simulated under (A) constant birth-death, (B) increasing speciation-rate, and (C) recent mass-extinction models and their jackknifed trees (dashed lines) at 90%, 80%, 70%, 60%, 50%, and 40% are plotted. As the tree moves further from complete, the density plot shifts left, as a result of a declining principal *λ*, and the shape of the spectral density becomes increasingly different from the original, notably by decreasing skewness. The mean and standard deviation of the Jensen-Shannon distance between each tree and its 100 jackknifed trees are shown in a barplot. The distance between trees increases linearly with incompleteness.

**Supplemental Figure 7:**
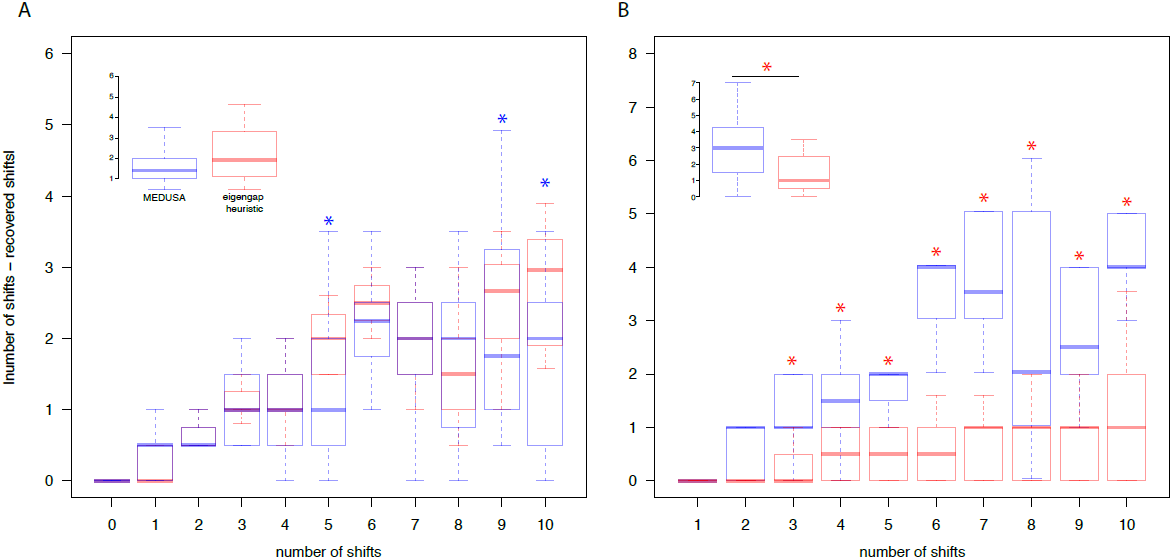
The reliability of the eigengap heuristic *versus* MEDUSA in recovering shifts in speciation rate and diversification pattern in simulated trees. The absolute deviation of shifts recovered by the eigengap (red) and MEDUSA (blue) from the known number of shifts for trees with 0-10 shifts in (A) speciation rate and (B) diversification pattern. In (A), only when five or ten shifts were simulated, MEDUSA performed significantly better (T > 2, P < 0.05) than the eigengap. (A, Inset) The average deviation across all trees is slightly lower for MEDUSA (2.89) than for the eigengap heuristic (3.10), although this is not significant (T = 1.88, P > 0.05). In (B), the eigengap heuristic outperformed MEDUSA for all trees with > 2 shifts and (B, inset) overall (T = 10.44, P ¡ 0.01). In (A) and (B), only eigengaps supported by *BIC* post-hoc analysis were computed in the means. Asterisks indicate a significantly lower deviation for MEDUSA (blue) or the eigengap heuristic (red).

**Supplemental Figure 8:**
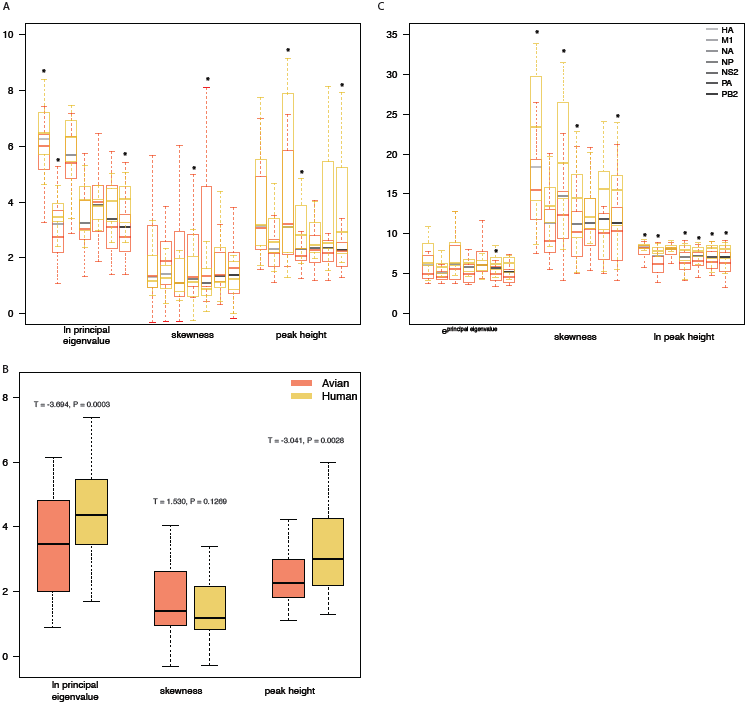
Spectral density profile summary statistics across viral strains and hosts. Boxplot of avian (red) and human (gold) strains calculated from standard (A) and normalized (B) graph Laplacians. Grey bars indicate across-host means; asterisks denote significant differences at P < 0.05. (C) Mean differences between hosts across all strains calculated from standard graph Laplacians.

**Supplemental Figure 9:**
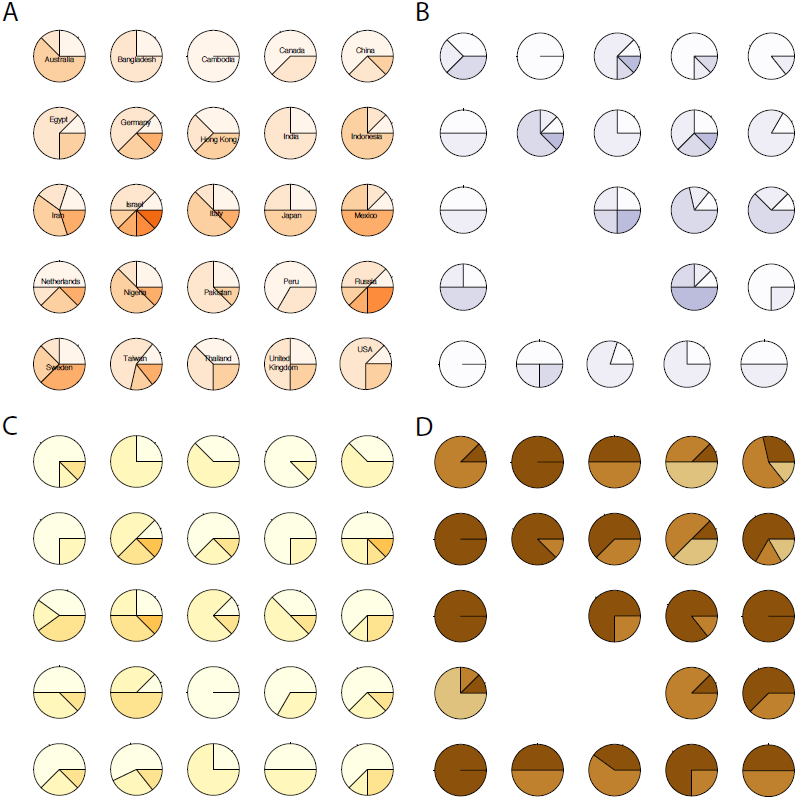
The cluster assignments for strains by country of origin. Six clusters were found based on k-medoid clustering on the standard spectral density profiles of all strains (P < 0.05). The distribution across those clusters for strains sampled from 25 countries are shown for avian (A) and human (B) hosts. Four clusters were found using the normalized profiles (P < 0.05) and the distributions are shown for each country for avian (C) and human (D) hosts. Only countries with all seven strains sampled are represented.

